# Functional microRNA-Targeting Drug Discovery by Graph-Based Deep Learning

**DOI:** 10.1101/2023.01.13.524005

**Authors:** Arash Keshavarzi Arshadi, Milad Salem, Heather Karner, Kristle Garcia, Abolfazl Arab, Jiann Shiun Yuan, Hani Goodarzi

**Affiliations:** Burnett School of Biomedical Sciences, College of Medicine, University of Central Florida, Orlando, FL, USA; Department of Computer Engineering, University of Central Florida, Orlando, FL, USA; Department of Urology, University of California, San Francisco, San Francisco, CA, USA; Helen Diller Family Comprehensive Cancer Center, University of California, San Francisco, San Francisco, CA, USA; Department of Biochemistry and Biophysics, University of California, San Francisco, San Francisco, CA, USA; Bakar Computational Health Sciences Institute, University of California, San Francisco, San Francisco, CA, USA

## Abstract

MicroRNAs are recognized as key drivers in many cancers, but targeting them with small molecules remains a challenge. We present RiboStrike, a deep learning framework that identifies small molecules against specific microRNAs. To demonstrate its capabilities, we applied it to microRNA-21 (miR-21), a known driver of breast cancer. To ensure the selected molecules only targeted miR-21 and not other microRNAs, we also performed a counter-screen against DICER, an enzyme involved in microRNA biogenesis. Additionally, we used auxiliary models to evaluate toxicity and select the best candidates. Using datasets from various sources, we screened a pool of nine million molecules and identified eight, three of which showed anti-miR-21 activity in both reporter assays and RNA sequencing experiments. One of these was also tested in mouse models of breast cancer, resulting in a significant reduction of lung metastases. These results demonstrate RiboStrike’s ability to effectively screen for microRNA-targeting compounds in cancer.

## Introduction

As a class of short non-coding RNAs, microRNAs (miRNAs) are among the essential regulators of cellular homeostasis. They oversee gene expression and regulate protein synthesis across many target regulons. The dysregulation of these post-transcriptional regulatory programs has been shown to contribute to tumor formation and progression [1], as part of oncogenic and/or tumor suppressive pathways.

Since microRNAs regulate a variety of gene regulatory programs, they play an important role in the emergence of various oncogenic hallmarks, such as metastasis [2], angiogenesis [3], and resistance to apoptosis [4]. There is recent evidence that they may contribute to the suppression of the immune response within the tumor microenvironment as well. It has also been established that microRNAs are involved in drug resistance in multiple cancers [5]. Among miRNAs, miR-21 has been one of the most well-studied drivers of oncogenesis [6]. miR-21 dysregulation has been implicated in ovary, gallbladder, colorectal, pancreatic, and many other cancers.

Much effort has been dedicated to targeting microRNAs to combat various tumors. This is largely achieved through the synthesis and delivery of antisense oligonucleotides (ASOs). For example, FDA researchers developed ASOs that are capable of targeting the long terminal repeats of ERV-9, which is known to be involved in multiple types of cancer, including breast, liver, and prostate tumors [7]. However, delivering ASOs to cells can be difficult due to their poor permeability, and short ASOs may also bind to other RNAs with similar sequences, leading to unintended effects. To overcome these limitations, some researchers have focused on developing small molecules that bind and inhibit miRNAs [8, 9]. Unlike ASOs, small molecules are typically easier to formulate, and have better bioavailability, making them more suitable for drug development. In recent years, researchers have made progress in developing small molecules that target various types of RNAs, both coding and non-coding. Risdiplam, for example, is a small molecule in early stages of discovery for its ability to treat triple-negative breast cancer and Alport syndrome by inducing the degradation of miR-21 via RNase L recruitment. As another example, Risdiplam is a FDA approved drug for its ability to treat Spinal Muscular Atrophy (SMA) by targeting SMN2 pre-mRNA exon 7– intron junction [8].

It has been difficult, however, to structurally inhibit the activity of RNAs with small molecules, especially microRNAs. Unlike proteins, they typically have a dynamic structure [10]. Since microRNAs lack the canonical pockets found in druggable proteins, conventional docking approaches have not been successful for microRNAs [11]. Furthermore, structurally binding a small molecule to microRNAs does not necessarily impair their functionality [12]. Another approach to targeting microRNAs is to inhibit their upstream regulators, but these regulators are largely unknown, making them difficult to target. Despite significant research efforts, microRNAs are still considered to be largely undruggable, and a platform that can specifically target the activity of different microRNAs has been a long-sought goal in the field.

In this study, we use machine learning to develop a small molecule drug discovery platform called RiboStrike, which aims to inhibit microRNA activity rather than disrupting their structure. Our approach is built on the capabilities of advanced deep learning architectures to learn representations from molecular data and discover hidden patterns in an abstract and non-linear manner. As shown in Figure 1, Graph Convolutional Neural Networks (GCNNs) [13] are used to aid the virtual screening of small molecules against miR-21 solely relying on the graphbased input. We used Multitask Learning to learn patterns from multiple data sources including large publicly available assays from PubChem [14]. Furthermore, multiple modules were added to the pipeline to improve its performance and utility, such as uncertainty prediction, auxiliary modeling (e.g. toxicity prediction), and molecular diversification, to help prioritize molecules for wet lab follow up and validation. RiboStrike selected eight candidate compounds from a pool of more than 9 million molecules in the ZINC database [15]. We then performed a battery of experiments, in cell culture and in mice, to functionally validate the anti-miR21 activity of the hit compounds identified by Ribostrike. Taken together our results indicate that RiboStrike can identify candidate inhibitory molecules without the need for the structural information of the target. Additionally, in this work, we have introduced a prediction-based algorithm for recommending input datasets for multitask learning in order to improve the performance of the modeling for the discovery of molecules that inhibit miR-21 activity.

**Figure 1:**
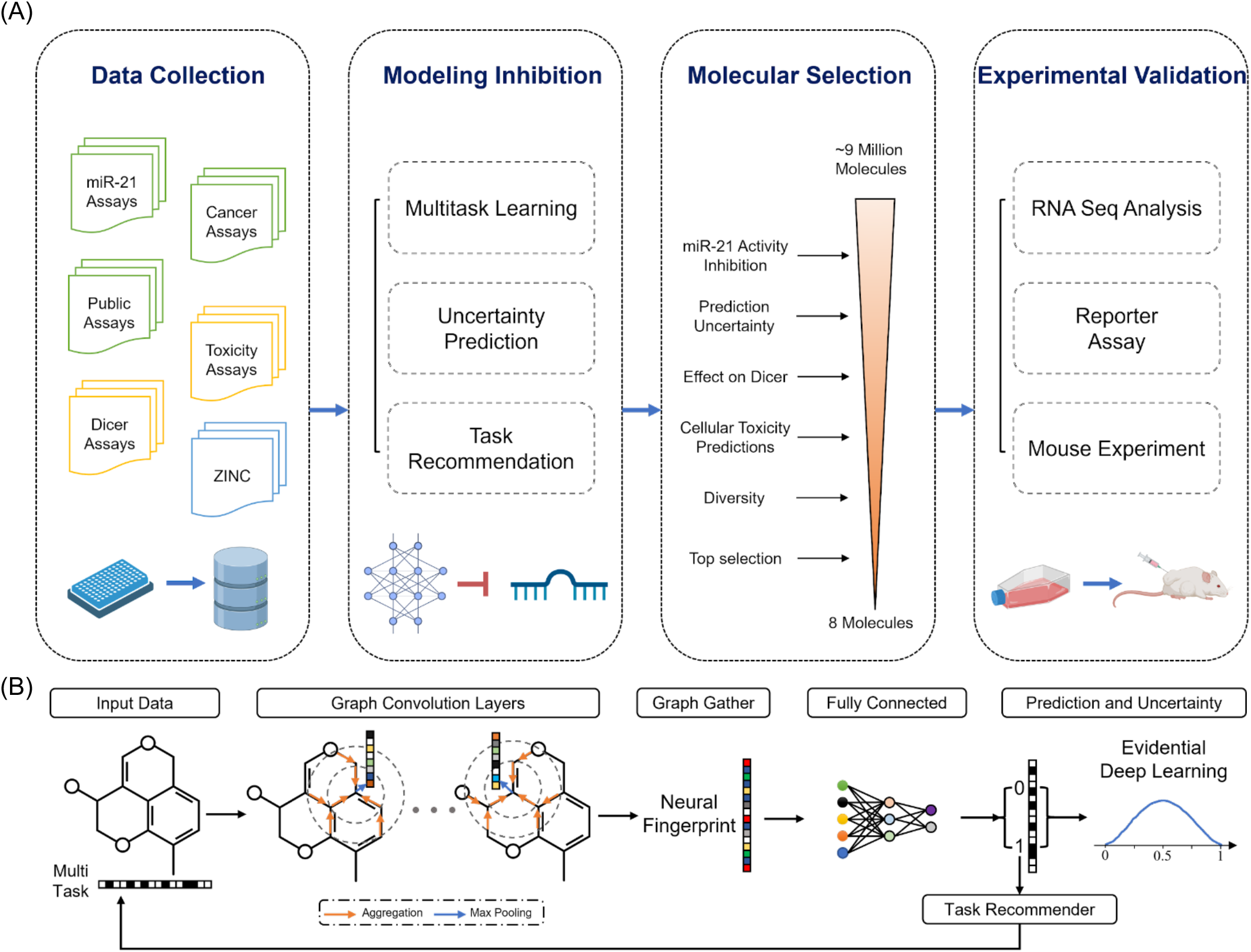
Overview of the RiboStrike pipeline. **(A)** Stages of discovery, from input molecular data and deep learning techniques, to candidate selection and experimental validation. **(B)** Computational pipeline and the flow of data within the GCNN network.

## Results

### Aggregating Data for miR-21, Cancer, and Off-Target Interactions

We used three different data modalities to train RiboStrike for optimized hit detection. First, we used datasets relevant to the virtual screening task to train models that predict the effect of a particular small molecule on the activity of miR-21. In this category, we combined nearly 14K inhibitors of miR-21 activity (measured by a reporter assay in a 315K-compound library) [16], assays designed to identify cancer-fighting candidates (Cell-based and biochemical assays), and the PCBA dataset (a collection of overlapping PubChem assays [17]). Combining all these datasets resulted in a total of 139 tasks, however, in order to create a more focused training dataset that is optimal for multitask learning, we used a prediction-based task recommender algorithm to narrow the combination of the three mentioned datasets down to seven tasks. Additionally, we also took advantage of a dataset that measures DICER inhibition. DICER is one of the main processing enzymes for microRNAs and its inclusion allowed us to counter select against DICER activity and identify miR-21 specific inhibitors as opposed to systemic inhibitors of miRNA biogenesis. This model, alongside a model trained on general cellular toxicity, guides the pipeline’s prediction to maximize on-target efficacy and specificity and filter overtly toxic compounds. Finally, we used large scale libraries of drug-like *in silico* compounds as inference datasets to find candidates for further screening. These datasets include ZINC [15] for its diverse and large collection of molecules and Asinex [18] for its novel selection of molecules. We have listed a summary of all datasets used for training, inference, and filtering in this research in Table 1.

**Table 1.** Summary of the datasets. used for virtual screening (VS), Auxiliary Modeling (AUX), and molecule selection (MS), as well as the number of tasks (assays), number of molecules, and the percentage of active molecules within each dataset.

| Dataset | Type | Tasks | # Molec. | Active (%) |
| --- | --- | --- | --- | --- |
| miR-21 [16] | VS | 1 | 315.16K | 4.29 |
| Cancer | VS | 48 | 535.45K | 2.54 |
| PCBA [17] | VS | 128 | 439K | 1.39 |
| Combined | VS | 139 | 540.42K | 2.39 |
| Recommended | VS | 7 | 448.87K | 4.53 |
| Toxicity [19] | AUX | 58 | 8.36K | 18.73 |
| Toxicity Recommended | AUX | 5 | 8.36K | 19.64 |
| DICER [20] | AUX | 1 | 46.72K | 6.05 |
| ZINC [15] | MS | - | 9.20M | - |
| Asinex [18] | MS | - | 3652 | - |
| FDA Approved [21] | MS | - | 157 | - |
| SAR Sample [22] | MS | - | 37 | - |

### Computational Pipeline - Training Graph Convolutional Neural Networks on Small Molecules

In learning from molecular data, RiboStrike utilizes GCNNs as its primary modeling approach. Graph-based models are well-suited for handling small molecules since nodes represent atoms within molecules and edges represent bonds between them [13, 23]. In this graph representation, each node contains features that describe the atom and its properties. Through the use of these features during training, abstract representations of the nodes can be formed by applying graph convolution which can then be pooled into a representation for the given molecule through the graph gathering layer. We and others have used GCNNs to effectively learn the relationship between input molecules and their physicochemical properties [24], their impact on a given target’s function [25] and virtual screening [26]. Other models such as pretrained transformers (e.g. ChemBERTa [27]) which rely on masked token prediction from SMILES sequence or tensor field networks [28] and extract features from coordinate clouds, were also considered for this work. However, GCNNs were ultimately chosen due to their simplicity, smaller size, and training efficiency (empirical comparison of performance to ChemBERTa can be found in Supplementary Materials). In this study, we have also taken advantage of evidential deep learning [29] to provide an estimate of the uncertainty of a prediction given an input molecule. As shown in Figure 1, we trained GCNN models to predict miR-21 activity suppression, DICER inhibition, and toxicity profiles of each input molecule, given their canonical SMILES (Simplified molecular-input line-entry system). We have illustrated our pipeline for the flow of data in this study in Figure 1B.

### Task Recommendation - Tailoring the Training Dataset for miR-21

Despite the large dataset created by combining all assays, the high number of tasks often degrades the model’s performance. A particular problem may arise during multitask learning and Stochastic Gradient Descent (SGD) when training on multiple tasks may reduce the performance on one or more of the tasks (e.g. some tasks differ from others regarding the gradient direction), which results in a less efficient training process [30]. As a result, we implemented a novel prediction-based recommendation algorithm to select a subset of tasks from the dataset and to create a smaller dataset after the multitask model has been trained on all tasks. Using this algorithm, sub-models with similar predictions to the target task (for example, miR-21) are identified, and the top-ranking tasks relating to these sub-models are selected as recommendations. Figure 2 provides an overview of the scores for all submodels on the target task as well as the threshold line. Using our recommender technique, we identified seven tasks that scored higher than the threshold of the mean plus two standard deviations (The tasks are identified in the Supplementary materials Table S2).

**Figure 2:**
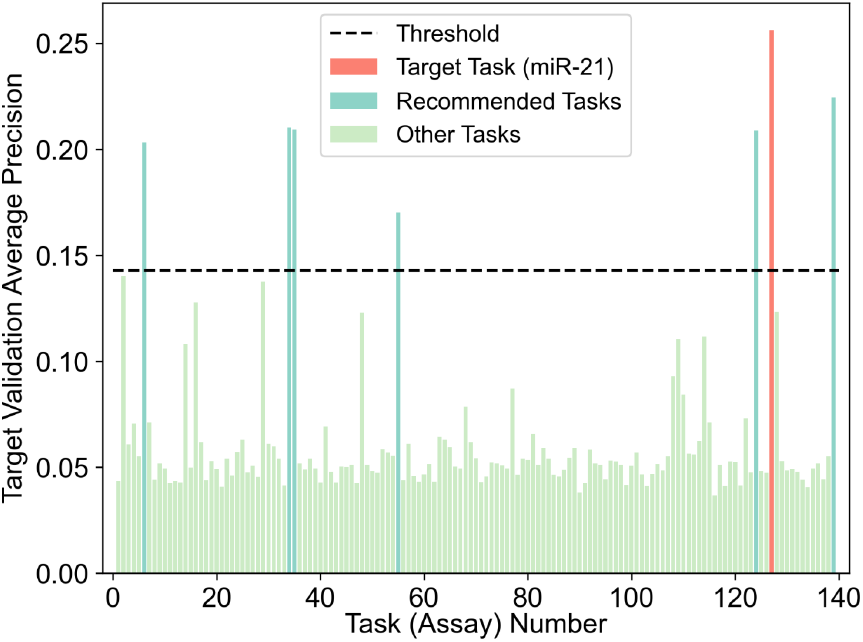
The Average Precision (AP) score of different submodels for the task recommender algorithm. The tasks above the threshold line make predictions matching the miR-21 ground truth to a higher degree than the rest of the tasks. These tasks are the recommended tasks and are selected for training a new model.

As expected, the counter screen assay for miR-21 activity [31], which was an assay to detect false positives in the main screen by measuring the activator of the firefly luciferase, is positioned as one of the recommended tasks. This is intuitive due to this counterscreen’s direct association with the main task and the importance of molecular patterns and activity labels contained in this dataset. Interestingly, the remaining recommended tasks were not directly related to miR-21, and were included solely based on the observation that the output of their trained sub-model is similar to that of the miR-21 sub-model. In other words, the models trained on these recommended tasks perform relatively better in predicting the effect on miR-21 activity than the rest of the submodels, making their respective training data suitable for use in the recommended model.

### Virtual Screening Results - Task Recommendation Offers the Best Performance

Following identification of the optimized tasks and implementation of all training scenarios, we compared a variety of modeling techniques to ascertain which dataset and training regimen resulted in the best-performing model. The results on the test set, 10% of the original miR-21 dataset which was held out and isolated throughout model training, were calculated and summarized in Table 2. The confusion matrices for three different training scenarios are shown in Figure 3.

**Figure 3:**
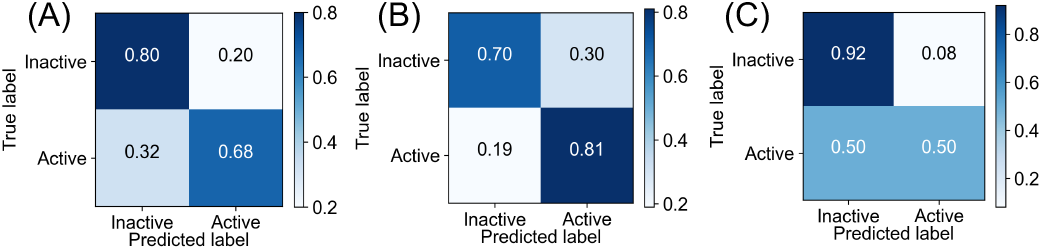
Confusion matrix for models trained using different learning methods. **(A)** single task, **(B)** multitask for all tasks, and **(C)** multitask for recommended tasks.

**Table 2.** Performance of different models on the test set of the miR-21 dataset. Models have different types of training data, yet all contain the same miR-21 task for fair comparison.

| Model | # Task | ROC-AUC | Precision | AP |
| --- | --- | --- | --- | --- |
| Single Task | 1 | 0.8170 | 12.52 | 23.03 |
| Multitask - Cancer | 48 | 0.8239 | 11.10 | 20.62 |
| Multitask - All | 139 | 0.8325 | 10.14 | 20.43 |
| Random Selected Tasks | 7 | 0.8027 | 18.11 | 21.04 |
| Recommended Tasks | 7 | 0.8305 | 20.93 | **24.72** |

As was expected, the prediction-based task recommender algorithm resulted in the highest-performing model using the recommended tasks, compared to random tasks, all tasks, or single tasks, as the model trained on the recommended tasks achieves the highest average precision score. Since this model has a lower tendency to predict molecules as active, it suffers from a loss in recall. It is evident, however, from the precision score in Table 2 and the confusion matrix in Figure 3C, that molecules that are predicted to be active are more likely to be true positives, which is a desirable behavior for virtual screening models.

### Evaluation of the Ribostrike model on held-out datasets

To independently verify the performance of our best-performing model, we used the Structural Activity Relationship sample dataset [22] which includes 37 molecules derived from two inhibitors of miR-21 that can overcome chemotherapy resistance in cancers such as renal cell carcinoma. Our virtual screening model successfully classified 33 out of 37 molecules as “active”, closely matching the results of the related study [22], demonstrating the potential for this model in the context of SAR scenarios. For the four misclassified molecules, our model was uncertain (with uncertainty of 100% for all four misclassified cases). This is likely because, for the most part, these molecules were in the last iteration of SAR, and were therefore significantly altered. Consequently, these molecules are outside the familiarity zone of our model’s training set, resulting in an uncertain prediction for the model.

### Auxiliary Models for Selecting Molecule Candidate: DICER Inhibition Modeling Results

MicroRNAs are transcribed and matured through a predefined pathway. DICER is the main processing enzyme for microRNA biogenesis and inhibiting its activity will reduce the activity of all microRNAs and not just miR-21. To ensure that our model is resistant to this possibility, we added a counter screening against DICER to ensure miR-21 specificity of the model [32]. For this, we trained a specialized model on data from an assay regarding DICER-mediated maturation of pre-microRNA [32] to predict inhibitory activities against the DICER as a way to identify and avoid unwanted inhibitory effects. The performance of this model is shown in Table 3.

**Table 3.** Performance of toxicity and DICER inhibition models on their corresponding test sets. The models belonging to the toxicity category are comparable.

| Model | # Task | ROC-AUC | Precision | AP |
| --- | --- | --- | --- | --- |
| DICER Single Task | 1 | 0.6631 | 12.22 | **12.53** |
| Tox Multitask | 58 | 0.7408 | 41.13 | 38.41 |
| Tox Recommended | 5 | 0.7483 | 38.27 | **38.73** |

### Auxiliary Models for Selecting Molecule Candidate: Toxicity Modeling Results

It is imperative that candidates for drug development are non-toxic and with as few off-target interactions as possible. As a way to identify any unwanted inhibitory effects on selected targets that could lead to cell toxicity, we trained two toxicity models. In order to determine molecular toxicity against cell viability, we used data from HepG2, a liver cell line that is used as a standard model. Moreover, we trained additional auxiliary GCNN models on the Tox21 dataset [19], which includes 58 different toxicity tasks, with the purpose of filtering compounds that may be broadly toxic to cells. Overall, the molecules with few toxicity predictions (out of 58) and the lowest uncertainty on HepG2 toxicity prediction pass satisfy this filter. The performance of these two models is described in Table 3. More detail on the result of toxicity and DICER inhibition models are included in the Supplementary materials.

According to Table 3, the model trained on the recommended tasks is capable of predicting HepG2 toxicity with a higher AP. In summary, this model is used to predict HepG2 toxicity and uncertainty, whereas the multitask model is used to perform the rest of the toxicity tasks.

### Selecting Diverse Candidates by Clustering the Learned Small Molecule Embeddings

Once the virtual screening model and the auxiliary models have been trained, they can be used to screen novel molecules for their potential as drug candidates. Given its large and diverse collection of drug-like and synthesizable compounds, we used the ZINC15 library for our virtual screening; however, we also included compounds from the Asinex library, which represents an out-of-distribution collection and allows us to assess the generalizability of our model. We used our trained model to screen these datasets for suitable molecules that are most likely to specifically inhibit miR-21 activity with fewer potential side effects and least likely to cause overt toxicity. We first used our multitask virtual screening model trained on the seven recommended tasks to predict miR-21 activity inhibition across 9 million molecules *in silico*.

To obtain a more complete understanding of how diverse the inference molecules are, we performed unsupervised analysis and clustering of the molecular embeddings learned by the model. This analysis also enabled us to select molecules from various regions of the molecular space. To accomplish this, the embedding features of the trained model were extracted, projected into a 2D UMAP space for visualization, and clustered using the KMeans algorithm. By separating molecules into clusters, different regions of the molecular space can be accessed to select more diverse molecules for follow up and testing (Figure 4).

**Figure 4:**
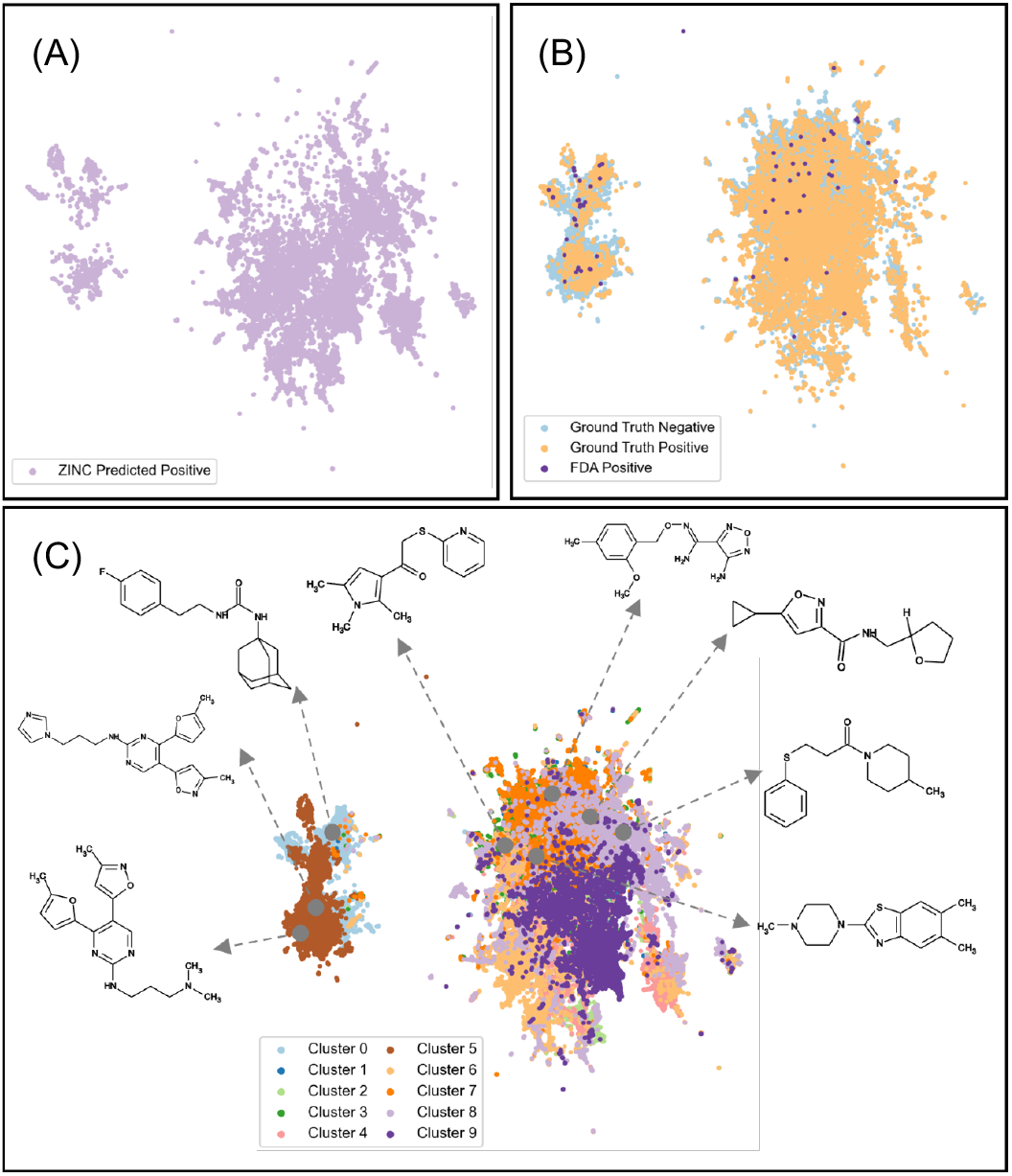
**UMAP of the inner features of the model for the inference and training data for (A)** ZINC data predicted to be active, **(B)** the training data and the positive molecules from the FDA Approved set, and **(C)** the 10 clusters applied to this space for the inference sets (ZINC and Asinex) and the final selected molecules.

### Assessment of the Learned Small Molecules Embeddings and selection of Hit Compounds for Further Validation

The positive and negative training data distributions in Figure 4B have a substantial overlap, which highlights the challenge in virtual screening for miR-21. It is also evident from Figure 4A that the molecules from the ZINC database that have been predicted to be positive cover most of the projected space, demonstrating that the model is not learning singular molecular features. It is also visible that FDA-approved drugs [21] are concentrated in certain clusters, which likely makes the selection of hit molecules from their vicinity favorable. By clustering the hit molecules into ten groups across the embedding space, we ensured that our selected hit compounds were sampled from varying clusters and therefore retained molecular diversity.

Using clustering as the basis for the selection of molecules, those hit compounds that are predicted to affect the activity of miR-21 with high confidence were selected and virtually screened against DICER inhibition and cellular toxicity. To conclude, we selected a total of ten molecules across the two inference datasets (ZINC and Asinex) specifically from embedding clusters where FDA-approved molecules are well-represented. We selected these molecules from different clusters to ensure diversity. We also required them to have low uncertainty in predicted inhibition of miR-21’s activity, with few or no toxic activity predictions, and without predicted activity against DICER. Overall, six molecules were selected from the ZINC dataset and four from Asinex. Of these, we successfully acquired eight molecules, six from ZINC and two from Asinex. We ensured that none of these molecules had been previously studied in this context and by any large represented new chemical entities. The selected molecules are identified by their IDs and SMILES in Supplementary Materials Table S3.

### Gene expression profiling to measure miR-21 activity in response to treatment with the selected hit compounds

Since miR-21 is a post-transcriptional regulator of RNA stability, reducing its activity results in an increase in the RNA levels of its target regulon. To experimentally verify the anti-miR-21 activity of our selected compounds, we used an RNA sequencing strategy amenable to scalable gene expression profiling, namely QuantseqPOOL. For experimental testing, we used MDA-MB-231 cells, an established model of triple negative breast cancer metastasis that is known to be driven by miR-21 [33]. Inhibition of miR-21 in these cells should significantly reduce their metastatic potential. We first used CellTiter-Glo to calculate the IC-20 for each of our compounds, to ensure a regimen in which the key cellular processes are not impacted by each treatment. We then performed QuantseqPOOL on MDA-MB-231 cells treated at IC-20 for 72 hours in biological replicates. We also included DMSO-treated control samples. For positive control, we used an established anti-miR-21 ASO and included a non-targeting ASO as control. Upon measuring the gene expression changes induced by each compound, we asked whether they caused a systematic effect on the expression of miR-21 target RNAs. We used the set of RNAs that are annotated as miR-21 targets (based on Targetscan [34]) to perform gene-set enrichment analyses (Figure 5A). As expected, in the miR-21 ASO samples, we observed a significant enrichment of miR-21 targets among genes that were up-regulated upon treatment. From the eight selected compounds, two show miR-21 target up-regulation similar to that of the ASO; and a total of five show some activity (with a confidence score of over 85%). Based on these results, our model has a hit rate of 62.5%, higher than the *in silico* predicted precision score of 20.93% for our model.

**Figure 5:**
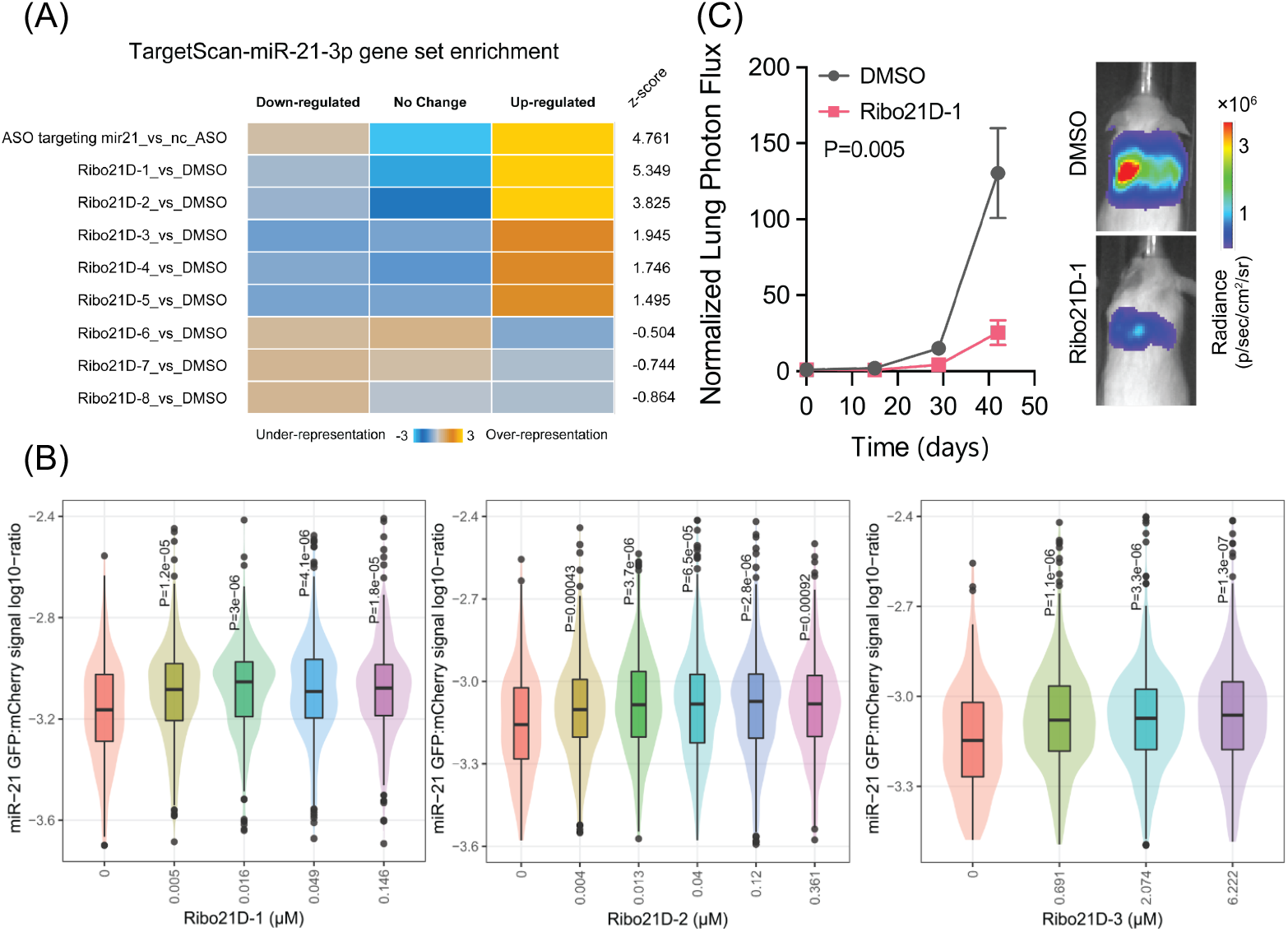
Experimental Validation Results. **(A)** gene set enrichment analysis to sort candidate molecule based on upregulating effect on miR-21-3p target genes. **(B)** Reporter assay results. **(C)** Mouse experiment results.

### Reporter assays for targeted measurement of miR-21 activity in dose-response assays

For the five compounds showing anti-miR-21 activity based on RNA-seq data, we also used a reporter assay for an independent confirmation at multiple doses. For this, we designed a GFP reporter harboring two miR-21 recognition sites in its 3’ untranslated region (3’UTR). This construct also drives the expression of an mCherry gene, which serves as an endogenous control for the construct [35]. We used miR-21 ASOs and cytometry to confirm that inhibiting miR-21 resulted in a significant increase in GFP expression. We then performed this experiment for each compound separately as well. As shown in Figure 5B, three of the five compounds also showed significant dose-dependent activity against miR-21. Taken together, our platform showed an experimentally confirmed hit rate of 37.5% for entirely new chemical entities that were predicted *in silico*.

### Measurement of anti-metastatic activity in xenograft mouse models

As mentioned earlier, miR-21 is a driver of metastasis in breast cancer. Therefore, we expect the inhibition of this miRNA to result in lower metastatic potential in breast cancer. To confirm this, we treated MDA-MB-231 breast cancer cells for 48 hours with the most promising candidate molecule, MCULE-9082109585. We then injected these cells, along with mock-treated controls, into immunocompromised Nod-Scid Gamma (NSG) mice via their tail-veins. We used *in vivo* imaging to monitor the colonization and growth of cancer cells in the lungs of mice over a period of roughly 40 days. As shown in Figure 5C, we observed that, consistent with its anti-miR21 activity, pre-treating breast cancer cells with MCULE-9082109585 resulted in a significant reduction in their lung colonization capacity. Therefore, not only this compound reduced miR-21 activity, as measured by RNA-seq and reporter assays, but also reduced metastatic lung colonization in xenografted mice. Together, our findings establish the utility of Ribostrike as an effective platform for discovering compounds against microRNAs.

## Discussion

In light of the dynamic structure and small size of miRNAs, the discovery of inhibitory small molecule against them presents a number of challenges. This is further compounded by the fact that microRNA ligands do not necessarily interfere with the function and activity of microRNAs. Considering this, we propose discovering candidate hits that instead focus on targeting microRNA activity. The goal of this study was to implement a virtual screening platform that leverages deep learning to enable the selection of early hit candidates out of a large collection of diverse molecules. Multiple methods were implemented in order to ensure the practicality of the computational methods, as well as rigorous experimental characterization, resulting in the validation of multiple compounds *in vitro* and for our top hit also *in vivo*. The first step involved the use of deep learning models to train on a large number of small molecule datasets to learn the chemical language underlying miR-21 activity. We also introduced a new task recommendation technique, which identified the optimal configuration for combining datasets to maximize training potential. The third methodology involved calculating uncertainty for all predictions made by the models, which enabled the ranking of molecules in spite of their binary nature. To finalize, the internal features of the model were used to represent molecules during inference, and clustering of these embedding features allowed the selection of a diverse set of molecules for experimental testing. In total, the RiboStrike platform identified multiple hit candidates that were subsequently confirmed, demonstrating the advantage of using graph-based deep learning to identify hidden patterns of molecular hits against the activity of microRNAs without the need for sequence reading or structural information. It is likely that these compounds target molecular pathways upstream of microRNAs that regulate their processing, lifespan, and function. Therefore, once identified and validated, these hits can be used as toy compounds to identify their direct targets, e.g. using CRISPRi screen with microRNA reporter expression as read out [36]. The identification of these direct targets are not only required for additional genetic and biochemical validations of their impact on the activity of specific microRNAs, but this knowledge provides an avenue for further optimization of our tool compounds using traditional SAR and medicinal chemistry. Therefore, this approach may provide us with a better understanding of the mechanisms by which microRNAs are regulated through the discovery of crucial unknown players upstream.

## Acknowledgements

We thank Dr. Kevan Shokat for reviewing the selected molecules and advising on the selection process. H.G. is an Era of Hope Scholar, and supported by the following grants: R01CA240984, R01CA244634, and W81XWH-20-1-0541.

## Author contributions

A.K.A collected the data, performed the molecular selection and helped with the implementation of the computational pipeline. M.S. implemented the code and the computational pipeline. H.K. and K.G. performed the in-culture experimental measurements. A.A. analyzed the RNA-seq data. H.G. and J.S.Y. supervised the research. A.K.A, M.S., and H.G. wrote the manuscript.

## Materials and Methods

### Data Sources

Three categories of data are used throughout the RiboStrike pipeline: virtual screening datasets, off-target interaction datasets, and inference datasets, which are explored further in the following subsections. Each dataset is preprocessed to contain canonical SMILES as well as appropriate binary labels for the activity of molecules, the details of this process can be found in Supplementary Materials.

### Virtual Screening Training Datasets

The datasets used to train the main virtual screening model or to help its performance via multitask learning are as follows:

- miR-21 Data: Primarily, an HTS dataset from miR-21 inhibition screening as a target is used in this study, a data repository originating from National Center for Advancing Translational Sciences (NCATS) and deposited in PubChem (ID: AID 2289 [16]). The aim of this assay had been the discovery of molecules with an inhibitory effect on miR-21 to finally induce cell apoptosis and tumor suppression. They used a cell-based firefly luciferase reporter gene assay optimized for qHTS.
- Cancer-Related Data: To assist the multitask training process, different cancer-related assays are collected from PubChem and the PCBA dataset. These assays include 20 tasks directly from the PubChem database, and 38 cancer-related tasks from the PCBA dataset [17].
- PCBA dataset: PubChem BioAssay (PCBA) is a collection of datasets aggregated from PubChem consisting of the biological activities of small molecules generated by high-throughput screening. In this work, a subsection of PCBA with 128 bioassays is used with over 400 thousand molecules, similar to the previous benchmarking methods [17]. This dataset was selected due to its size, high number of tasks, and high molecular overlap with the miR-21 dataset. Due to these features, this dataset can be combined with the miR-21 dataset to create a large non-sparse training set for multitask learning.

Two further datasets are created from the mentioned dataset; the “combined” dataset from aggregating all data points, and the “recommended” dataset from algorithmically selecting tasks from the combined dataset using the task recommendation approach (will be discussed in Prediction-Based Task Recommendation section).

### Off-Target Interactions Datasets: Toxicity and DICER Inhibition

To increase the probability of any predicted bioactive molecule to be a drug candidate, the toxicity properties and the effect of these molecules on the DICER’s function are predicted. Therefore, two datasets related to these subjects are used for training in this work:

- Tox-21 (Toxicity in the 21st century): the most popular source available for the analysis of the toxicity of molecules [19]. This dataset consists of 58 distinct tasks each tested on an important protein target of humans. Candidate drugs that interact with those targets are likely to cause side effects and cellular toxicity. Therefore, we used 58 tasks from tox21 to filter out molecules that would show side effects in future steps.
- DICER dataset: The DICER protein plays an important role in the maturation of several RNAs, not just miR-21. Therefore, if a candidate drug decreases the activity of the miR-21 by inhibiting the DICER protein, it would show significant side effects downstream. The dataset for this task is taken from PubChem data source of AID 1347074 [20]. This dataset was created using click chemistry, and was selected due to the fact that this assay identified inhibitors of DICER based on pre-miR-21.

### Inference Data - Sources for Drug Candidate Selections

There are multiple datasets used in this work for the drug candidate selection:

- ZINC: ZINC database is a diverse molecular library typically used for virtual screening [15]. It contains millions of molecules that are indexed for search and properties. We used the drug-like subset (MW: 250 to 500, LogP: −1 to 5) of the ZINC15 database, consisting of 9 million molecules that had 3D representations, standard reactivity, reference pH, a charge of −2 to +2, and were in stock for purchase [37].
- Asinex: Asinex is a molecular library vendor that sells varieties of different classes of molecules including macrocycles, alpha helix mimetics and peptidomimetics. This vendor also offers a small molecule library specifically designed to target RNA [18], which is a suitable candidate for this project. Moreover, the candidates in this set are different from the training set, allowing for testing the model’s generalizability.
- FDA Approved: This dataset was taken from a study [21] which found 86 molecules that were suppressor of miR-21 activity from a pool of 696 compounds from the Bioactive Compound Library and 262 compounds from FDA-approved Drug Library. In this work, these molecules are deleted from the mentioned inference datasets, to avoid selection of previously discovered molecules. Moreover, these molecules will be used to assist the selection of the final molecules by identifying the favorable clusters as described in the molecule selection section.
- SAR Sample: This dataset originates from a study [22] which takes two molecules that are known inhibitors of miR-21, and uses Structural Activity Relationship (SAR) to optimize these molecules and assist in overcoming chemoresistance in Renal Cell Carcinoma. Overall, 37 molecules are tested and shown to be active, which can create a small validation set for this work.

### Graph Convolutional Neural Network Training

In recent years GCNNs have proven to be helpful in learning representations from small molecules and modeling tasks such as virtual screening [25], molecular property prediction [24], and drug-target interaction [38, 39]. This success is owed to two facts, firstly small molecules are inherently similar to graphs, with atoms represented as nodes and bonds represented as edges, making GCNNs suitable tools for handling this data type. Secondly, the feature extraction in the GCNN model, which is inspired by traditional circular fingerprint extraction from molecules, results in useful and often superior inner features due to the automatic representation learning aspect of deep learning [13, 40].

In this work, the GCNN implementation from the DeepChem library is used [41]. In this model, the molecules are converted to graphs and atoms, then featurized to include features such as atom type, number of directly bonded neighbors, implicit valence, formal charge, and hybridization type. Isomeric information is also added to the features in the form of a vector with length of three (whether chirality is possible, right-hand, and left-hand). The hyper-parameters of this model as well as the length of the training are found through hyperparameter optimization in a grid search manner. (More on hyper-parameter optimization can be found in the Supplementary Materials)

### Evaluation Metric

The metric used for evaluation in this work is the average precision score (AP). This metric computes the area under the precision-recall curve and was chosen due its fairness towards imbalanced datasets, where the positive label discovery is of importance. This is the case with most virtual screening tasks, where the number of active molecules is often much lower than the number of inactive molecules, resulting in highly imbalanced datasets. Moreover, discovery of these active candidates is of utmost importance in an early drug discovery pipeline since these candidates will be passed on to the next steps of the drug discovery process. Therefore, average precision score is favored in this work for comparison of models, different architectures, or different epochs during training. The results are also reported for accuracy, recall, precision, and the area under Receiver Operator Curve (ROC-AUC).

### Optimizing the Tasks for Multitask Learning

Multitask learning has proven to be beneficial in many instances via providing multiple tasks for the model to simultaneously learn from, with the hope that the learned representations for these tasks benefit from being shared within the same model. However, this is not the case in all scenarios and in some cases negative transfer occurs, where multitask learning hurts the performance of a given task when compared to single task learning [30]. In this work, to address the problem of negative transfer, the novel method of “prediction-based task recommendation” is proposed, which narrows the number of tasks selected for multitask learning via recommending a few tasks in an algorithmic manner.

### Prediction-Based Task Recommendation

To begin the process of task recommendation and selection of fewer training tasks, a multitask learning model is trained on all available tasks. After training, one target task is selected (e.g. miR-21 dataset) and inference is performed on the validation set of this target task. Since the model is trained on multiple tasks, it will have multiple predictions for each input molecule, each assigned to one input task. Given *N* total training tasks and *M* molecules in the target task’s validation set, the predictions of the model will then have a shape of *M×N*, with each row representing the output of the sub-model assigned to one input task, denoted by *Output_i_*. After this output is calculated, each *Task_i_* is scored using the scoring metric in Equation 1.

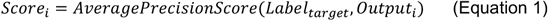

In which *Label_target_* denotes the ground truth labels for the target task. As it can be seen from this equation, the labels are kept constant on the target set while the submodel changes, which is the main difference between this method and simple inference. Using this scoring mechanism, the predictions of different sub-models are compared to the ground truth labels of the target task, with the sub-models that have more similar predictions to the target labels having a higher score. Through the identification of sub-models with similar predictions, this approach identifies their corresponding training tasks and selects the highest-scoring tasks for training. The recommended tasks are selected via applying a threshold of mean plus two standard deviations to the scores. This threshold is arbitrary and can be replaced with a simple selection of top K scores. The recommended tasks are then passed on as training data to the hyper-parameter optimization and training step for the final model. This recommendation process is repeated for the toxicity model as well, with the target task of HepG2.

### Molecule Selection: Inference and Uncertainty Prediction

After different categories of models are trained, the final models are used to predict the properties of the molecules from the inference datasets. All training data, as well as the FDA approved data [21], is removed from the inference datasets to avoid selection of redundant or previously discovered molecules. During the inference process, a binary label is predicted for each given molecule, reflecting its predicted effect on miR-21’s function. Having only binary labels to distinguish between different molecules is problematic, since molecules with the same predictions (e.g. active) become indistinguishable and no ranking can be assigned to the molecules for further drug candidate selection.

In order to overcome the selection challenges, an uncertainty prediction method is applied on the last layer of the model. To do so, evidential deep learning is used [29] which applies a Dirichlet distribution on the class probabilities and computes uncertainty for each prediction. This uncertainty score ranges from zero to one, with lower scores demonstrating more certain predictions. With this uncertainty score, the predictions become distinguishable and the molecules that are predicted to be active with low uncertainty become desirable.

### Molecule Selection: Neural Fingerprints Clustering

After the uncertainty is predicted, one challenge is faced which is the lack of diversity in the top selected molecules with low uncertainty. The reason for this phenomenon is that similar molecules result in similar predictions and uncertainty, and the most certain predicted molecules are similar to each other and they populate the top of the certainty ranking list. This creates a problem in the further stages and specifically in vitro screening, where diversity among the candidates is needed to increase the chance of activity against different targets.

To enforce diversity within the selected molecules and create variety within the final selection, the molecules are clustered, and a few molecules are selected from some of the clusters. To do so, the neural fingerprints of the molecules are extracted from the Graph Gather layer of the trained model. This fingerprint is the inner features of the model for an input molecule and is a numerical vector that can meaningfully represent this molecule. Neural fingerprint clustering allows the molecules belonging to different clusters to be both structurally different, and exist in different locations within the feature space of the trained network. After the features are extracted, KMeans (K=10) clustering is applied to the features, then visualized using 2-dimensional Uniform Manifold Approximation (UMAP) projection, resulting in 10 clusters formed from the inference molecules.

### Molecule Selection: Five Criteria for Final Selection

After the molecules are clustered and uncertainty and bioactivity is predicted for all of them, five different criteria were checked for the final molecules to be selected:

1. Potency as miR-21 activity inhibitor: The selected molecules should be predicted to inhibit the function of miR-21.
2. Certainty: The molecules that had the least uncertainty in each cluster were considered.
3. Diversity: Molecules should belong to different clusters in regard to the clustering of the neural fingerprint. Clusters that include more of the FDA approved molecules are more likely to be selected from.
4. Pass majority of toxicity tests: The selected molecules are more likely to be predicted as non-toxic in most of the toxicity tests with low uncertainty for specifically the HepG2 test.
5. Low chance of inhibiting the DICER: The selected molecules are more likely to be predicted to not affect the function of the DICER with low uncertainty.

Following these criteria, the inference molecules are first narrowed down to those predicted to be active with high certainty, then filtered via selecting top molecules from each 10 clusters. Afterwards, the final molecules are selected from this list with consideration of toxicity and DICER activity and their uncertainties. In the end, 8 molecules are selected from the inference datasets (ZINC and Asinex), and progress to the in vitro screening stage.

### Cell culture

The MDA-MB-231 (MDA-parental, ATCC HTB-26) human breast cancer cell line; its highly metastatic derivative, MDA-LM2 [42]; its triple reporter version, MDA-MB-231tr; and HEK293T cells (ATCC CRL-3216) were cultured in Dulbecco’s modified Eagle medium supplemented with 10% fetal bovine serum (FBS), penicillin, streptomycin, and amphotericin B. Cells were all incubated at 37oC at 5% CO2 in a humidified incubator.

### Cell Titer-Glo Cell Viability Assay

MDA-MB-231 cells were seeded at 1k/well in white opaque 96-well plates (Corning 3917) and treated with serial dilutions of drug candidates ranging from 3.2pM to 1mM for 72 hours in triplicate. Cell viability was measured using CellTiter-Glo 2.0 Assay (Promega G9243) to find IC20 concentrations.

### High-throughput sequencing data generation

MDA-MB-231 cells were seeded at 3k/well in clear 96-well plates and treated with drug candidates at IC20 for 72 hours in duplicate. Controls were transfected with oligo inhibitors targeting either mir21 or non-targeting controls in duplicate. Per well, 0.5 uL 100 uM inhibitor, 100 uL OptiMEM (Thermo Fisher Scientific 31985088), and 2.5 uL Lipofectamine 2000 (Thermo Fisher Scientific 11668019) were incubated for 20 minutes at room temperature before being added to cells and incubated at 37c for 24 hours. RNA was extracted using the Quick-RNA 96 kit (Zymo Research R1052) and concentrations were determined using Nanodrop. Libraries were prepared using the QuantSeq-Pool Sample-Barcoded 3’mRNA-Seq kit (Lexogen 139) with 10 ng input RNA. Libraries were sequenced using NovaSeq 6000 SP (100 cycles).

### RNA-seq data analysis

The QuantSeq Pool data demultiplexed and preprocessed using an implementation of the pipeline provided by Lexogen – https://github.com/Lexogen-Tools/quantseqpool_analysis. The outputs of this step are gene level counts for all samples. The raw counts matrix used for differential expression analysis using DESeq2 package [43]. The log fold changes from multiple differential expression comparisons used gene set analysis (Figure 5A).

Finally, we aimed to sort molecules based on their systematic effect on expression of miR-21 target mRNAs through gene set enrichment analysis. Thus, we downloaded TargetScan prediction of hsa-mir-21 3p target genes from miRBase dataset (Accession: MI0000077). Then, we use a modified version of iPAGE [44] (we call it onePAGE from here) in which you can perform the gene set enrichment analysis powered by mutual information evaluation and statistical tests for a single gene set. The onePAGE analysis here reports enrichment of miR-21 target gene list in 3 bins of log fold change values; from left to right 0) lowest bin of log2FC, i.e., downregulated, 1) log2FC around zero, i.e., no expression change. 2) highest bin of log2FC, i.e., upregulated. After running this analysis for differential expression log2FC of all drugs and control conditions, we sorted drugs based on resulted z-score with keeping miR-21 vs negative control (nc) at the top (Figure 5A). From this step, we selected the top 5 drugs for further evaluation.

All scripts for preprocessing, differential expression, onePAGE enrichment analysis are accessible in this GitHub repository – https://github.com/goodarzilab/targeting-miR-21-RNA-seq.

### Generation of MDA EGFPmiR-21 reporter cell line

The vector backbone for the reporter plasmids was generated from a vector with a bi-directional CMV promoter driven lentiviral reporter, expressing eGFP and ΔlnGFR. This vector was a gift from David Erle [45]. The ΔlnGFR ORF in the vector was replaced by a PuroR-T2A-mCherry fusion using Gibson assembly as previously described [46]. In order to generate the eGFPmiR-21 reporter plasmid two miR-21 binding sites were added to the end of eGFP using the NEBuilder HiFi DNA Assembly Cloning Kit. MDA-MD-231 cells were engineered to stably express the reporter plasmid using lentiviral delivery of the vector. (hsa-miR-21-5p, TCAACATCAGTCTGATAAGCTA, Sequences for reporter cell line generation were obtained from miRBase)

### FLOW cytometry analysis of miR-21 reporter cell line

MDA EGFPmiR-21 reporter cell lines were seeded at a density of 2×10^4 cells per well of a 96-well plate. Cells were treated for 3 days with serial dilutions of the 6 most promising drugs (dilutions determined by IC20 curves). Additionally, anti-miR controls (anti-mi21 and nontargeting) were also transiently transfected into cells using Lipofectamine 2000 (ThermoFisher) according to the manufacturer’s protocol. Cells were collected after 3 days and fluorescence output was measured on a BD FACSCelesta flow cytometer.

### Animal Studies

All animal studies were performed according to IACUC guidelines (IACUC approval number AN194337-01G). Age-matched female NOD scid gamma mice (Jackson Labs, 005557) were used for metastatic lung colonization assays. These assays were performed with MDA-MB-231 cells which were seeded at 1.5×10^5 cells in 2 wells of a 6-well plate on Day 1. After 24 hours 0.1uM MCULE-9082109585 and 0.5% of DMSO were added dropwise to one well each. After 48 hours, MCULE-9082109585 and 0.5% of DMSO media was removed and the cells were prepped for tail vein injections. Cells were resuspended in 2ml PBS and each mouse received 5×10^4 cells/100 ul of PBS. Metastasis was measured by bioluminescent imaging (IVIS), and histology performed by hematoxylin and eosin (H&E) staining of lung tissue sections.

## Supplementary Materials

### Molecular Data Preprocessing

After gathering multiple datasets for bioactivity classification, off-target interaction prediction, and inference, the molecules are represented in SMILES format. Each entry is then transformed to be canon with isomeric information included, using the rdkit library in Python (v3.6). Afterwards, the data is cleaned and missing, or duplicate entries are removed. The molecules from the toxicity dataset undergo one more preprocessing step, during which the molecules were desalted, and inorganic molecules were removed. This is since many molecules exist in the toxicity dataset that are salted, are redundant or contain inorganic atoms. Since these molecules are not relevant to this work’s screening, they were removed.

### Molecular Data Preprocessing: Label Assignment

In order to assign bioactivity labels to each molecule from the PubChem datasets, the two columns of “outcome activity” and “phenotype activity” were examined. In most assays, the datapoints with “Activator” phenotype were removed, since inhibitors alone were the desirable positive label for this work. In two cases (AIDs 504466 and 624202) the inhibitors were removed instead, since inhibitor and activators had the opposite meaning compared to other assays. For the remaining molecules, the outcome activity column is used to assign bioactivity labels, with ‘Active’ and ‘Inconclusive’ outcomes being labeled as 1, and ‘Inactive’ and ‘Unspecified’ outcomes being labeled as 0.

### Molecular Data Preprocessing: Data Splitting

The combined dataset, toxicity dataset, and DICER dataset are all split into training (80%), validation (10%), and test sets (10%) using DeepChem, based on the molecular scaffolds [47]. Splitting based on the scaffolds allows a larger distinction to exist between the molecules of each split, resulting in a more practical model for real-world applications where inference data is often from a different distribution than the training data. Moreover, splitting the combined dataset instead of each of the virtual screening datasets allows for the test dataset to remain constant throughout all types of training trials, resulting in a fair comparison between the models.

### Hyper-Parameter Optimization: Training

After the data is gathered and preprocessed to have binary labels and canonical SMILES, hyper-parameter optimization is performed to find the suitable model architecture for each training task. Using the validation set, different model architectures are evaluated for their performance. In a grid search manner, the number of graph convolutional layers, the size of each layer, the size of the feed-forward layer, the batch size, and the learning rate are altered and the best performing set of parameters on the validation set is taken to initiate the training. This hyper-parameter optimization is extended to the training epoch selection, where the epoch with the best performance on the validation set is chosen as the final model.

### Hyper-Parameter Optimization: Results

The results are shown in Table S1 for the virtual screening models. All models are trained for 150 epochs and the best epoch is identified using the AP score on the validation set.

As seen in Table S1, the hyper-parameter search arrives at similar optimum architectures for all datasets, with the model for the recommended task having a smaller batch size and training duration. This is intuitive since this training scenario has fewer tasks and a smaller training dataset.

**Table S1.**
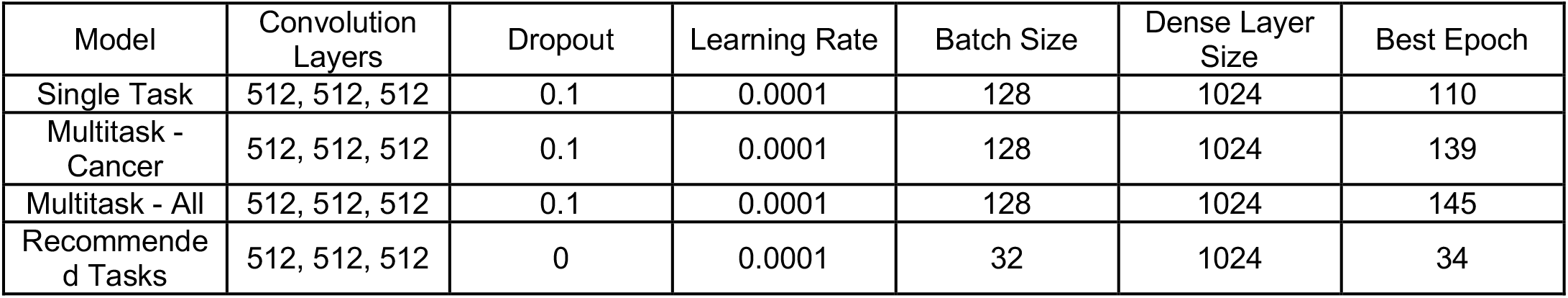
Optimum hyper-parameters for each training scenario of the virtual screening models. Best epochs are found through early stopping monitoring the validation set.

| Model | Convolution Layers | Dropout | Learning Rate | Batch Size | Dense Layer Size | Best Epoch |
| --- | --- | --- | --- | --- | --- | --- |
| Single Task | 512, 512, 512 | 0.1 | 0.0001 | 128 | 1024 | 110 |
| Multitask - Cancer | 512, 512, 512 | 0.1 | 0.0001 | 128 | 1024 | 139 |
| Multitask - All | 512, 512, 512 | 0.1 | 0.0001 | 128 | 1024 | 145 |
| Recommended Tasks | 512, 512, 512 | 0 | 0.0001 | 32 | 1024 | 34 |

#### Task Recommendation Results: Tailored Tasks for miR-21

The task recommendation algorithm is implemented to find tasks within the input dataset that when used for data augmentation, would result in a better performance for our target task of miR-21. Table S2 lists the tasks that were chosen and recommended.

**Table S2.** The recommended and optimized tasks for multitask learning for miR-21 virtual screening. Each assay is represented by its Assay Identifier (AID) in PubChem.

| Recommended Task Name | Importance |
| --- | --- |
| AID-1458: qHTS Assay for Enhancers of SMN2 Splice Variant Expression | Other/Unrelated |
| AID-485297: qHTS Assay for Rab9 Promoter Activators | Other/Unrelated |
| AID-485313: qHTS Assay for NPC1 Promoter Activators | Other/Unrelated |
| AID-588342: qHTS profiling assay for firefly luciferase inhibitor/activator using purified enzyme and Km concentrations of substrates (counterscreen for miR-21 project) | Counter Screen |
| AID-504466: qHTS screen for small molecules that induce genotoxicity in human embryonic kidney (HEK293T) cells expressing luciferase-tagged ELG1 | Other/Unrelated |
| AID-624202: qHTS Assay to Identify Small Molecule Activators of BRCA1 Expression | Other/Unrelated |
| AID-2289: qHTS Assay for Modulators of miRNAs and/or Inhibitors of miR-21 | Main Screen |

### Toxicity and DICER Inhibition Modeling Results

For the case of DICER inhibition prediction, a single task model is trained on the available data. On the other hand, the toxicity dataset contains multiple assays, which requires the same training pipeline (multitask learning as well as task recommendation) as the miR-21 virtual screening model to be reimplemented. In this process after a model is trained on all of the toxicity tasks, the prediction-based recommendation algorithm returns 5 tasks that would assist the performance of the model on the HepG2 target task.

### Comparison to ChemBERTa: Language Modeling for Small Molecules

In recent years, language models have gained popularity in modeling text within the natural language processing field as well as other sequence-based data such as proteins and even small molecules within the biology and chemistry fields. These models have the capability to learn from unlabeled sequences solely by relying on a mechanism called “masked token prediction”, where certain words or “tokens” (e.g. amino acids, nucleotide bases, or atoms within a SMILES string) are omitted and the model is asked to predict the missing tokens. By doing so, the model can learn the grammar and syntax of the sequences, and with this knowledge, create meaningful representations for these sequences. ChemBERTa [27] is a language model trained on 77 million SMILES strings from PubChem, which deploys masked token prediction and has a distinctive feature space to distinguish small molecules given their SMILES.

To assess the performance of our GCNN-based model, we compare it to ChemBERTa. The pre-trained model was taken from HuggingFace repository and modified to be fit for a classification task by adding one fully connected layer to the end of the model (with 32 neurons). The weights of the base model were frozen during training, except for the Layer Norm parameters to perform selective finetuning [48]. The model was trained for 20 epochs to predict the binary labels of the miR-21 task. The results are shown in Table S2.

**Table S2.** Performance of this work’s best model compared to ChemBERTa on the test set of the miR-21 dataset. Models are trained and tested on the same datasets.

| Model | Method | ROC-AUC | Accuracy | Recall | Precision | AP |
| --- | --- | --- | --- | --- | --- | --- |
| RiboStrike | multitask learning | 0.8305 | 90.39 | 49.75 | 20.93 | 24.72 |
| ChemBERTa | masked language modeling | 0.7547 | 88.57 | 36.40 | 14.22 | 11.89 |

As it can be seen from Table S2, the approach proposed in this work can empirically outperform ChemBERTa, with the help of multitask learning as well as task recommendation. One reason for ChemBERTa’s lower performance can be the resolution of the fingerprints it offers. As we have seen from Figure 4B, the current dataset has massive similarity between the positive and negative ground truth data points. This similarity would necessitate high resolution fingerprints to be leveraged to discriminate between the data points of different classes, while ChemBERTa’s inner representations are general and lack this resolution. Overall, the comparison made here is for using ChemBERTa as a feature extractor as it is the prevalent approach, it is noteworthy to mention that the pipeline of RiboStrike can be model-agnostic and the principals of multitask learning and task recommendation can be extended to any model, even pre-trained language models such as ChemBERTa.

### Molecular IDs

The name, IDs, and the SMILES for the molecules selected in this work for experimental validation are mentioned in Table S3.

**Table S3.**
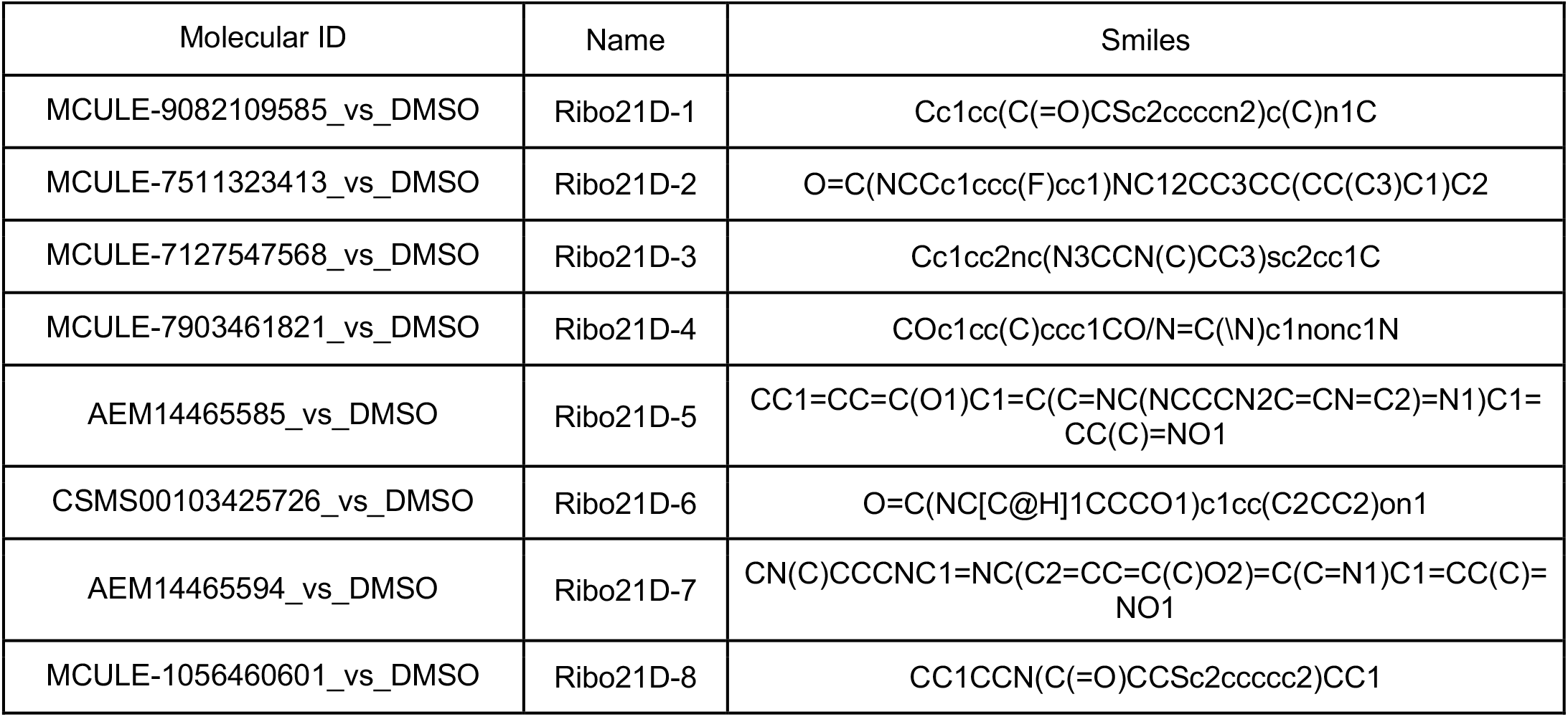
The selected molecules by the RiboStrike pipeline. Molecules within the text are referred to by their assigned names and can be found through their molecular IDs.

